# Rapid environmental DNA detection with isothermal amplification methods: a comparison with the real-time quantitative PCR in the natterjack toad (*Epidalea calamita*)

**DOI:** 10.64898/2026.11.06.764843

**Authors:** Abigaël Chieux, Alessia Guerrieri, Eva Bellemain, Frédéric Boyer, Thomas Merzi, Pierre Taberlet, Amaia Iribar, Lucie Zinger, Aurelie Bonin

**Author notes:** co-last authors (Amaia Iribar, Lucie Zinger, Aurelie Bonin).

## Abstract

Species detection relying on environmental DNA (eDNA) traditionally relies on Polymerase Chain Reaction (PCR). This amplification technique takes in general a minimum of one hour and requires in most cases access to specialized laboratories and equipment. Isothermal amplification techniques, originating from clinical research, are emerging as rapid and field-deployable alternative methods. Yet, whether their sensitivity, specificity, and cost/time-efficiency are comparable to the traditional real-time quantitative PCR (qPCR) remains to be assessed. In this work, we compare the performance of loop-mediated isothermal amplification (LAMP) and recombinase polymerase amplification (RPA) with qPCR to detect the natterjack toad (*Epidalea calamita)* from aquatic filtrations. This objective was achieved in three steps. First, we developed LAMP and RPA primers specific to the target species based on its mitochondrial genome. Second, we amplified DNA extracts from *E. calamita* and from tissues of closely related species using real-time fluorescence detection with LAMP, RPA, and qPCR commercially available protocols to evaluate and compare the sensitivity and specificity of the three methods. Finally, we validated all protocols on eDNA extracts recovered from water samples from ponds surveyed by field scientists. We show that RPA had a sensitivity equivalent to that of the qPCR, while LAMP sensitivity was ten times lower. All our assays amplified the species’ DNA in eDNA samples where the species was observed by field scientists. Through this work, we thus propose a robust methodology for implementing isothermal amplification assays for eDNA applications and show that RPA is a good alternative to qPCR for detecting conservation-priority species (i.e. protected or invasive species) when time and lab facilities are constrained.

## 1 Introduction

The ability to accurately detect the presence of a species in a given location is essential for biodiversity assessment and conservation purposes (IPBES 2019). It is key for tracking threatened taxa, to gain a deeper understanding of their spatial distribution, ecological interactions, population health, and the threats to their persistence. Such information is also crucial for managing ecological pressures, such as invasive species or pathogen outbreaks, and ultimately guides ecosystem management, ecological decisions and policymaking. Species recording is traditionally based on direct human detection through conventional field surveys including visual, acoustics or trapping methods (Sutherland 2006; Eymann et al. 2016; Wich and Piel 2021). While these conventional fieldwork methods are flexible and provide precious ecological insights, they are often constrained by high logistical cost, a reliance on scarce taxonomic expertise and difficulties to detect rare or elusive species (Fediajevaite et al. 2021). With the evolution of technologies, the last two decades have offered new opportunities for low-invasive, scalable, and time-efficient monitoring that complements direct field surveys. Among them, species detection based on environmental DNA (eDNA) infers the presence of species from the DNA traces they left in the environment (Taberlet et al. 2018). This approach unlocks the possibility to detect cryptic, rare or microscopic species that conventional field surveys may miss.

Traditionally, eDNA detection relies on one thermal cycling amplification method: the polymerase chain reaction (PCR) (Mullis et al. 1986). This process enables the exponential replication of short, taxonomically informative DNA fragments targeted by either generalists (multi-specific) or specific primers through repeated temperature-dependent cycles. Following the success of this gold standard, numerous variants of PCR protocols have been developed to enhance eDNA-based species detection sensitivity, specificity, and quantification. Examples include the widely used quantitative real-time PCR (qPCR; Langlois et al. 2021), as well as the increasingly employed digital PCR (dPCR; Capo et al. 2021) and more complex multiplex PCR (Jo et al. 2020; Lammers et al. 2024). Although PCR offers numerous advantages, it requires a costly bulky thermal cycling instrument with precise temperature control. This implies off-site analysis, which can result in significant delays to the obtaining of results and, ultimately, to the efficiency of ecosystem management. This critical point had led to the development of alternatives to PCR, called isothermal amplifications, which operate at a constant temperature and are thus suitable for point-of-care applications (Glökler et al. 2021).

Originally isothermal techniques were developed for clinical applications in the 1990s, typically for pathogen identification (Guatelli et al. 1990; Compton 1991; Walker et al. 1992), they were then democratized for SARS-CoV-2 diagnosis during the COVID-19 pandemic as fast, sensitive and low-cost alternatives to reverse transcription qPCR (Szobi et al. 2023; Chen et al. 2024). Among them, two are particularly well-suited for eDNA-based species detection: the Loop-Mediated Isothermal Amplification (LAMP; (Notomi et al. 2000) and the Recombinase Polymerase Amplification (RPA; (Piepenburg et al. 2006). They were developed for point-of-care applications, require minimal instrumentation, are relatively easy to implement and enable rapid and sensitive detection of target DNA. Briefly, LAMP operates at 60 to 65 °C and requires a single enzyme, a temperature-stable DNA polymerase derived from the bacterium *Bacillus stearothermophilus (Bst* DNA polymerase), capable of rapid DNA synthesis from a few template copies. It uses four to six specific primers that recognize six to eight regions of the target DNA. Amplification products form characteristic loops or hairpin-like structures of various sizes. LAMP amplicons can be read through a panel of detection methods including real-time fluorescence and colorimetry (Becherer et al. 2020). The main advantage of LAMP is the specificity and sensitivity of the amplification process, which is due to the large number of primers used. In contrast, RPA reaction works at a lower temperature, within a range of 37 to 42 °C, and involves three types of proteins: a recombinase, a strand-displacing DNA polymerase, and single-stranded DNA-binding proteins. Only two primers are required, which are 5 to 10 nucleotides longer than PCR ones. The recombinase locally opens the double-stranded DNA, allowing primers to bind and DNA synthesis to occur under constant temperature conditions. The strength of RPA is the low temperature that enables very fast and low-instrumented detection. Distinctive RPA detection methods exist, the most popular ones being real-time fluorescence detection, gel electrophoresis, and lateral flow strip detection (Li et al. 2018).

While these techniques were initially not developed for amplification of low concentrated and degraded DNA samples, they have already given encouraging results in the eDNA field for the past six years. For example, researchers have already successfully used both LAMP or RPA to detect fish (Hartle-Mougiou et al. 2024; Wei et al. 2023; Nakajima et al. 2024; Jothinarayanan et al. 2026), and insects (Peterson et al. 2023; Trujillo-González et al. 2022; Kim et al. 2026). Parasites were successfully detected through LAMP amplification (Jothinarayanan et al. 2026; Valencia et al. 2026), as were mollusks including clams and snails (Davis et al. 2020; Valencia et al. 2026), platypuses (Khodaparast et al. 2024), and invasive crayfish (Porco et al. 2022). RPA proved effective for jellyfish (Kim et al. 2024), salamanders (Ling et al. 2024), and zooplankton (Hu et al. 2024). These first proofs of concept emphasized the possibility to use isothermal amplification for eDNA detection and highlighted the speed of the amplification reaction, which takes less than 30 minutes for both LAMP and RPA versus minimum one hour for PCR. They also suggested that the detection sensitivity of these methods can sometimes surpass that of qPCR. Despite their potential, isothermal eDNA assays still face limitations that hold back their democratization. These include the primer design complexity to minimize non-specific products, as well as the limited number of reliable tools for predicting primer specificity *in silico* (Vasconcelos-Martins et al. 2026; Singh et al. 2026). Another challenge is the choice of the detection approach. Some of them can increase the overall risk of false positives because they imply tube opening after amplification, possibly releasing amplified DNA in the experimental area. In addition, reagent costs and limited accessibility due to the small number of commercial suppliers may further constrain their use (Yang et al. 2024; Singh et al. 2026). More importantly, no studies have directly compared LAMP and RPA specifically for eDNA detection to our knowledge.

Here, we assess whether LAMP and RPA isothermal amplifications could simplify eDNA-based species detection and therefore accelerate monitoring decisions. To determine whether these methods are applicable to eDNA, we evaluated their ability to reliably detect the presence of a species compared to qPCR. We use the natterjack toad, *Epidalea calamita* (*E. calamita* or *Bufo calamita* as basonym; Laurenti, 1768) as a case study. This toad species has specific characteristics that make it a particularly suitable model for developing specific eDNA detection methods. It reproduces in shallow, recent and temporary water sources in spring and summer. Its local abundance and the rapid metamorphosis of tadpoles in a restricted and low-competition environment should greatly facilitate the collection and detection of its DNA. The purpose of this study is hence threefold: (i) to develop LAMP and RPA isothermal amplification assays for the specific detection of *E. calamita* in aquatic eDNA samples, (ii) to compare their specificity and sensitivity to those of a qPCR assay already established for this species and (iii) to demonstrate their effectiveness as rapid and robust alternatives for DNA amplification in an eDNA context.

## 2 Materials and Methods

A general overview of our comparative methodological workflow is outlined in **Figure 1**. The specific details of each step as applied to this study are described in the text below.

**Figure 1.**
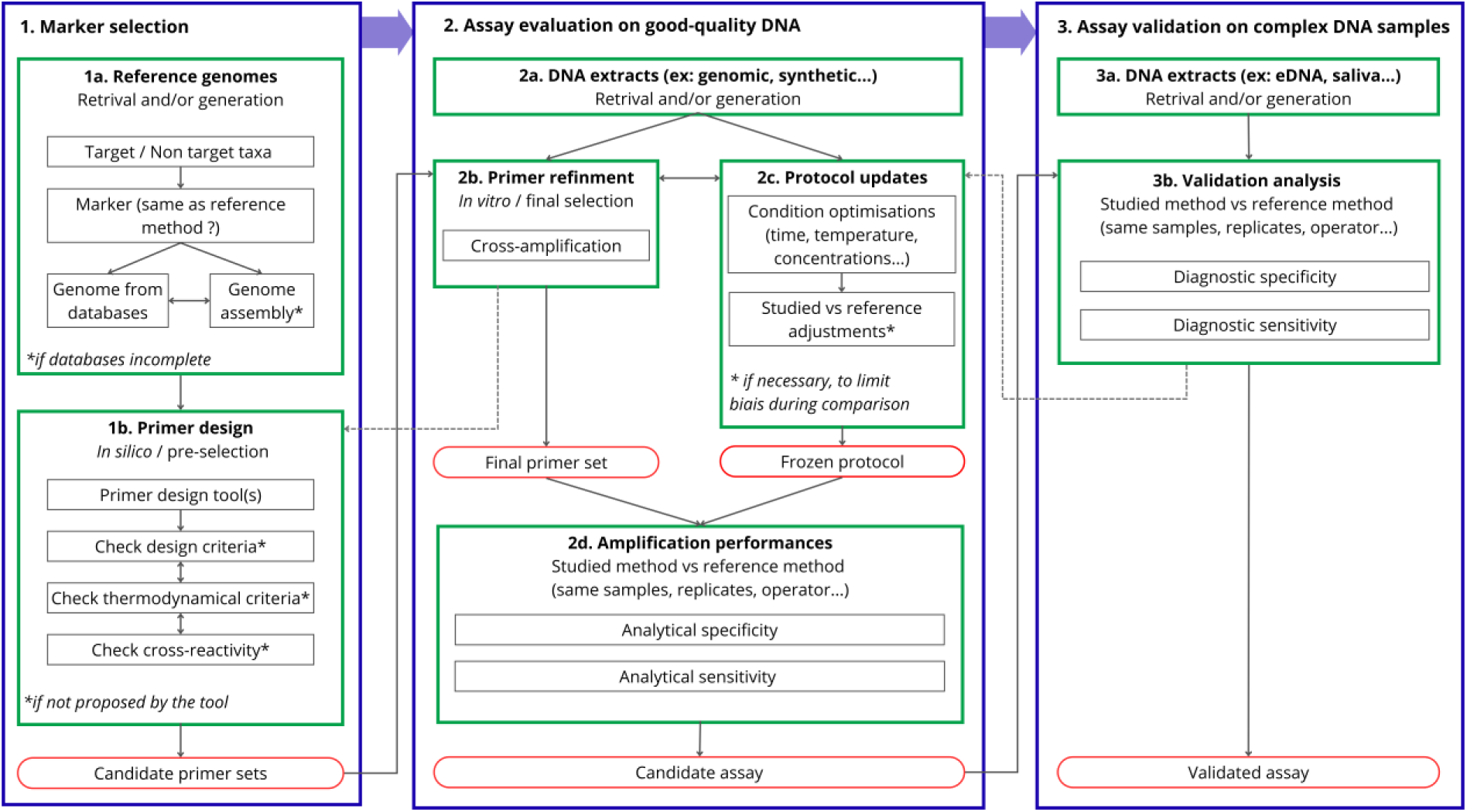
General methodology for the comparative study of DNA amplification methods on complex DNA samples. Blue boxes represent major steps, green boxes represent intermediate steps, black boxes represent criteria, and red and round boxes represent the final output of decision points. Solid arrows indicate workflow and dotted arrows indicate feedback.

### 2.1 Primers design

#### 2.1.1 LAMP primers design

LAMP primers were developed using the NEB® LAMP Primer Design software with default parameters (v1.4.2; **Table S1**). The reference mitochondrial genome of *E. calamita* (GenBank accession no: MT483697.1) was retrieved from the National Center for Biotechnology Information (NCBI) database in September 2024 and used as the input sequence (Sayers et al. 2025). The software generates candidate primer sets, both with and without loop primers. We retained only those with loop primers, as it is known to improve the reaction speed (Nagamine et al. 2002). Amongst the different candidate sets, we then selected those with the lowest extremity Gibbs free energy change ΔG° (i.e, large negative value, high annealing with template), as well as those with a higher primer dimer ΔG° (i.e low negative value, low annealing between primers) to ensure stable hybridization between primers and target DNA sequence during amplification. Ultimately, we identified five sets of six primers, each targeting different regions of the *E. calamita* mitochondrial genome, ready for subsequent *in vitro* testings (**Table S2**).

#### 2.1.2 RPA primers and fluorescent probe design

RPA primers were developed following the manufacturer’s recommendations described in the TwistAmp® Assay Design Manual (TwistDx, Abbott; [www.globalpointofcare.abbott]). We used the NCBI Primer-BLAST tool (Ye et al. 2012) which combines Primer 3 (Rozen and Skaletsky 1999) and BLAST (Altschul et al. 1990). While this tool is initially intended for PCR primer developments, the use of custom parameters allows for the design of RPA primers (Kellner et al. 2019). For these developments, we focused on two mitochondrial protein-coding genes, the cytochrome c oxidase subunit 1 (*COI*) and cytochrome B (*CYTB*), which are commonly used for PCR-based detection of amphibians. Two independent Primer-BLAST searches were conducted with custom parameters (**Table S3**). For each genomic region, *E. calamita* sequences, retrieved from the NCBI database in October 2024, were used as input (GenBank accession no: MT372456.1, *COI*; OQ130163.1, *CYTB)*. We then retained primer pairs with the least negative dimer and hairpin structures ΔG° values according to the OligoAnalyzer tool (Integrated DNA Technologies; [www.idtdna.com]) Then we checked the specificity of primers selected *in silico* with the Aliview alignment software (v1.28; Larsson 2014), using amphibians *CYTB* and *COI* sequences retrieved from NCBI. It consisted in 43 of the 45 amphibian species occurring in metropolitan France for the *COI* marker, and 40 species for the *CYTB* marker, with one to three sequences per species, corresponding to 70 *COI* sequences and 41 *CYTB* sequences in total. One human *COI* sequence and one human *CYTB* sequence were also included in the tests as human DNA is a likely source of contamination (**Table S4**). Three primer pairs per targeted genes were finally selected with slight shifts of a few bases amongst them (**Table S5**). We did so to facilitate the creation of a comparison matrix to obtain optimal primers combinations, as recommended in the TwistAmp Assay Design Manual. To enable real-time visualization of RPA amplicons, two exo probes were also designed for *CYTB* and *COI* primer pairs in accordance with the manual.

### 2.2 Assay evaluation on genomic DNA

#### 2.2.1 Tissue samples obtention

Tissue samples used in this study come from the CEFE - EPHE collection of the Montpellier Biogeography and Ecology of Vertebrates team (BEV collection). They include cellular material of 16 individuals collected at different stages of development and at various locations across Europe (**Table S6**). Six tissue samples were obtained from individuals of *E. calamita.* The remaining samples include a set of closely related species from the Bufonidae family with *Bufo bufo* (Linnaeus, 1758; n=2), *Bufo spinosus* (Daudin, 1803; n=3) *and Bufotes viridis* (Laurenti, 1768; n=2) as well as *Bombina variegata* (Linnaeus, 1761; n=2) *and Rana temporaria* (Linnaeus, 1758; n=1) from the order Anura. We extracted each sample using the NucleoSpin® Tissue kit (Machery-Nagel, Düren, Germany) with the total sample as starting quantity (< 25 mg) following the manufacturer’s protocol. A gel electrophoresis on 2% agarose gel was made to check the DNA quality, and DNA concentration was measured using a Qubit fluorometer (Invitrogen).

#### 2.2.2 Primer testing

LAMP and RPA primer testing in the laboratory was realized through preliminary amplification tests on *E. calamita* genomic DNA. All five LAMP sets underwent fluorescent amplifications as described below, and the final set was selected based on amplification performance results. For RPA, the primer screen was performed using end-point electrophoresis. The F1/R1, F1/R2, and F1/R3 combinations were evaluated for both target genes (**Table S5**). Primer combinations involving the F2 and F3 forward primers were not tested, as no further optimization was considered necessary. The primer pair selected was then used for fluorescence experiments with the exo probe following the protocol described in the section below entitled “*LAMP, RPA and qPCR assays*”. Minor variations in reaction volumes and reagent proportions for both LAMP and RPA protocols occurred between preliminary testing as reaction conditions were progressively refined during primer screening.

#### 2.2.3 LAMP, RPA and qPCR assays

To assess the effectiveness of LAMP and RPA amplifications, we compared them against qPCR amplifications. We chose to focus on fluorescence-based assays for LAMP and RPA as they allow for real-time monitoring of amplification and produce results that are easily comparable to qPCR. This approach also eliminates the need to open tubes after amplification, reducing the risk of cross-contamination and aerosol contamination of the experimental aera.

For LAMP assays, we used the WarmStart® LAMP Kit (DNA & RNA) for real-time fluorescence detection (New England Biolabs, Ipswich, MA). The LAMP primers were combined to prepare a 10X primer mix, with final concentrations of 16 μM for FIP and BIP, 2 μM for F3 and B3, and 4 μM for LF and LB. Each LAMP reaction was performed in a 20 µL reaction volume and contained 2 μL of template DNA, 10 µL of WarmStart® LAMP Master Mix (New England Biolabs, Ipswich, MA): 0.4 µL of 50X fluorescent dye, 2 µL of the 10X primer mix, and ultra-pure water to 20 µL. The reaction was incubated at 65°C for 45 min with fluorescence measurements recorded every 30 s on a Bio-Rad qPCR thermal cycler with a SYBR/FAM channel.

Each RPA reaction was prepared with the TwistAmp® Liquid exo kit (TwistDx Limited, Cambridge, UK). The final volume was 50 µL and the reaction mixture contained 2.1 µL of each primer (10 µM), 0.6 µL of a 10 µM exo probe, 25 µL of 2X reaction buffer, 1.8 µL of 50 mM dNTPs (Sigma-Aldrich, St. Louis, MO), 5.4 µL of ultrapure water, 2.5 µL of 20X core reaction mix and 1 µL of 50X Exo buffer. In the tube lid, we carefully added 2.5 µL of MgOAc and 2 µL of template DNA before agitation and centrifugation of the tubes, as the amplification starts only when the MgOAc is in contact with other reagents and DNA. Tubes were then rapidly put for incubation at 37 °C for 45 min on a Bio-Rad qPCR thermal cycler with a SYBR/FAM channel, with fluorescence measurements recorded every 30 s.

For qPCR amplifications, we used the primers from (Blom 2018) (Forward: TTACTTCACCAAGCAATATGACTATA, Reverse: TGTGTTGATGCTTAGATGCG, TaqMan® Probe: [6FAM]CACAATGTAACCTCCACGCTGAAAGAA[BHQ1]) targeting a 121-bp long 16S rDNA mitochondrial region. Each qPCR mix consisted of 2 µL of DNA template, 10 µL of AmpliTaq Gold® 360 Master Mix (ThermoFisher Scientific, Waltham, MA, USA), 0.16 µL of BSA, 2 µL of each primer (10 µM) and 1µL of probe (10 µM), as well ultra-pure to 20 µL. Amplification was carried out over 60 cycles. The qPCR program consisted of an initiation step at 95 °C for 10 min; a denaturation step at 95 °C for 30 sec; a hybridization step at 55 °C for 30 sec; and a final elongation step at 72 °C for 90 sec. Fluorescence was read with a Bio-Rad qPCR thermal cycler with the SYBR/FAM channel to read the dye.

#### 2.2.4 Assay specificity and sensitivity evaluation

Primer specificity was assessed by evaluating cross-reactivity through amplification tests using DNA extracted from natterjack toad tissue versus from closely related anuran species tissues described above (**Table S6**). To standardize the assays, we diluted all genomic DNA extracts in a concentration range between 0.1 and 0.9 ng/ul (i.e. the range of the least concentrated tissue extracts we obtained from the BEV collection. Each run of amplification was performed with three technical replicates for each DNA extract and the negative template control (NTC). We further assessed the detection sensitivity of each amplification method relative to qPCR. We subjected one genomic DNA extracted from *E. calamita* tissue mentioned above to seven serial dilutions from 0.1 ng/ul to 0.0001 pg/ul. Then each dilution was amplified in triplicate with each amplification method using the protocols described in the previous section “LAMP, RPA and qPCR assays”. Extraction and amplification negative controls were also amplified in triplicates in each run. The entire experiment was performed three independent times for each method to account for experimental variability and ensure the robustness of the sensitivity results.

### 2.3 Assay validation on eDNA samples

#### 2.3.1 Study sites

We collected eDNA water samples in two locations: Montmélian (Savoie, Auvergne-Rhône-Alpes, France) and Toulouse (Haute-Garonne, Occitanie, France). Based on local field scientists’ records, we selected sites which exhibit different levels of species’ presence (**Fig. 2**). In Montmélian, we selected two sites, one with confirmed presence (i.e., observation of eggs on the day of sampling, Site A; **Fig. 2**), and the other with confirmed absence serving as a negative control (Site B; **Fig. 2**). In Toulouse, we chose one site with high toad densities (i.e., observation of eggs the day of sampling and numerous individuals surveyed two weeks before, Site C; **Fig. 2**), one site with uncertain current presence (i.e., a reproduction site in 2015 where only one individual was observed along the path near the site in spring 2026, with no eggs or toad calls detected, Site D; **Fig. 2**), and one negative control site (Site E; **Fig. 2**).

**Figure 2.**
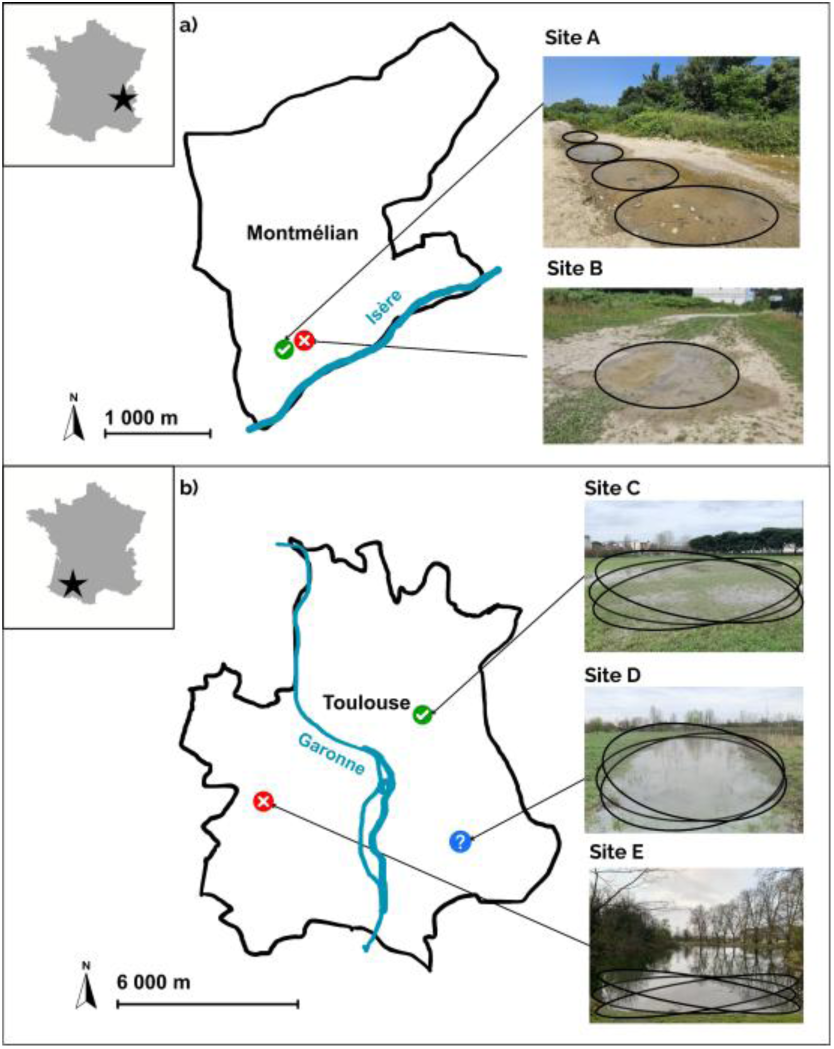
Environmental DNA sampling locations and sites. Territorial boundaries on the figure correspond to commune boundaries. The locations are marked with arrows and indicated by small colored circles on the map: sampling areas where species have been observed are shown in green (√), those where no species have been recorded (X) are shown in red, and the one where species have not been confirmed is in blue (?). Pictures of each site are on the left with black circles indicating the sampling zones of each composite sample. For Toulouse, circles correspond to biological replicates.

#### 2.3.2 eDNA sample collection

Samplings at Montmélian and Toulouse were carried out in June 2025 and March 2026, respectively. In both locations, and for each site, each sample was collected using a random point sampling plan, avoiding the bottom of the pools. We collected surface water using a ladle in Montmélian and a filtration pump (ARGALY, Montmélian, France) in Toulouse. Each sample therefore consisted of one liter (Montmélian) or two liters (Toulouse) collected in a Whirl-Pak**®** bag. We waited for the suspended matter to settle, and all the water was filtered using a 60 mL Luer Lock syringe connected to a filtration capsule (0.45 µm, 600 cm² filter; Waterra, Mississauga, Canada). After filtration, we emptied the filter by pushing air through the syringe a few times. Finally, we added 50 ml of Longmire preservation buffer to the capsule. The Longmire buffer was prepared according to the original protocol (Longmire et al. 1997) except that SDS was omitted and replaced by water. New gloves and a new 60 ml sterile syringe were used to collect each new sample. The ladle was also cleaned with bleach and rinsed with Evian water before moving on to the next sample. Overall, in Montmélian, four samples were collected in distinct areas adjacent to the egg masses observed at Site A, and one sample was collected at Site B (**Fig. 2**). In Toulouse, given the larger size of the water bodies, three samples were collected from the same area within each pool to obtain biological replicates. At Site E, access was limited to a single side of the pond so three biological replicates were collected from that section only (**Fig.2**).

#### 2.3.3 eDNA samples treatment

Back to the laboratory, capsules were agitated vigorously with a shaker in two directions for 15 sec each to release eDNA and cells from the filter membrane. Samples were transferred to 50 mL Falcon tubes, vortexed and centrifuged for 15 min at 15 000 g at 6°C. The supernatant was discarded, and DNA extraction was performed on the pellet. All samples were extracted using the NucleoSpin® eDNA Water kit (Macherey-Nagel, Düren, Germany), following the manufacturer’s protocol. An extraction negative control was included to monitor potential contaminants during this step. All DNA extracts were diluted at 1:5 and amplified through qPCR, LAMP, and RPA in triplicates using the same protocols as used for tissue amplifications (see section “LAMP, RPA and qPCR assays”). A positive control consisting of an *E. calamita* high-quality DNA extract, the one used for sensitivity tests, was included. Negative controls for each amplification method were also performed to monitor contamination at this step. To evaluate empirical specificity of the primers on eDNA samples, we also included in our tests eDNA water samples (Forêt de Marly, Saint-Nom-la-Bretèche, France), for which different European amphibian species have been detected by eDNA metabarcoding, but not *E. calamita* (**Table S7**).

### 2.4 Statistical analysis

All statistical analyses were performed using R (v.4.4.1; R Core team, 2024) and graphs were generated in Rstudio with the *ggplot2* R package (v3.5.2; (Wickham et al. 2007). All fluorescence results were obtained with the Bio-Rad CFX Maestro 2.3 program (v.5.3.022.1030; Bio-Rad Laboratories, Hercules, CA, USA). The single-baseline correction method was applied, and a fluorescence threshold was automatically determined by the software for each run (**Table S8**).

#### 2.4.1 Criteria for a positive amplification

For each amplification run, two metrics are extracted from the amplification curves by the software: (i) the quantification cycle (Cq), which is the cycle number at the intersection of the curve with the fluorescence threshold and (ii) the endpoint relative fluorescence units (EndRFU), representing the final fluorescence signal measured at the end of the reaction (**Fig. S1**). To identify which metric best discriminate positive from non-positive amplifications in our LAMP, RPA and qPCR assays, we used the amplifications results of the specificity tests (see section “*Assay specificity and sensitivity evaluation*”) and compared the distributions of EndRFU and Cq values between the target and non-target groups using non-parametric Mann–Whitney U tests. To facilitate comparison among methods, we converted Cq values into “time to threshold” (TTT) expressed in minutes. For qPCR, TTT for each reaction was calculated according to the following equation: TTTₓ = 10 + (2.5 × Cqₓ), where 10 min represents the initial denaturation and 2.5 min the duration per cycle. For isothermal amplifications, TTT was calculated as TTTₓ = Cqₓ/2 for each reaction, as the fluorescence was measured every 30 s. Based on these comparisons, EndRFU was identified as the metric providing the best discrimination between target and non-target amplifications and was therefore selected for determining a positive amplification for all analyses throughout the study. For this purpose, the run-specific EndRFU detection threshold automatically calculated by the CFX Maestro software was used. This threshold corresponds to the average EndRFU of the negative controls plus 10% of the range of EndRFU values across all samples within the run, using the software’s default setting. Samples with EndRFU values above this threshold are automatically assigned a positive amplification call by the software. The EndRFUs thresholds calculated for all runs throughout the study are reported in (**Table S8**).

#### 2.4.2 Sensitivity analyses

We modeled detection probability for each amplification method using a generalized linear model (GLM) with a binomial family and a logit link function (*lme4* package; v.1.1.37; (Bates et al. 2015), with the DNA extract concentration (log10-transformed and centered) and the amplification method as explanatory variables. The GLM model was previously adjusted. Detection probability corresponds to the probability of obtaining one, two or three positive amplifications among the three technical replicates. The qPCR method has been selected as the reference for the reasons explained above. Model validity was checked from residual analysis (DHARMa package; v.0.4.7; Hartig. 2026). We used *emmeans* package (v.2.0.1; Lenth et al. 2026) to perform pairwise comparisons to test differences in detection performance across methods. From this model, we also predicted the DNA concentration at which detection probability reach 95% (i.e. limit of detection at 95%, LOD95) for each amplification method (Bustin et al. 2009; Klymus et al. 2020). The corresponding 95% confidence interval was also estimated from the predicted values of the model.

## 3 Results

### 3.1 LAMP and RPA primers

Among the five candidates for LAMP (**Table S2**), one primer set was selected based on its amplification performance and consistency across replicates and independent trials using *E. calamita* tissue extracts. Our final choice resulted in a LAMP primer set that amplifies a 227 bp region of the mitochondrial genome of *E. calamita*. It targets the end of the *NADH dehydrogenase subunit II (ND2)* gene and three transfer RNA genes, trnW, trnA and trnN (**Table 1**). For RPA, candidate primer pairs targeting the mitochondrial *CYTB* and *COI* genes were screened. Primer combinations were tested sequentially, and screening was stopped once a primer pair showed satisfactory amplification performance. The selected primers target a 157 bp section of the *CYTB* mitochondrial gene (**Table 1**).

**Table 1.**
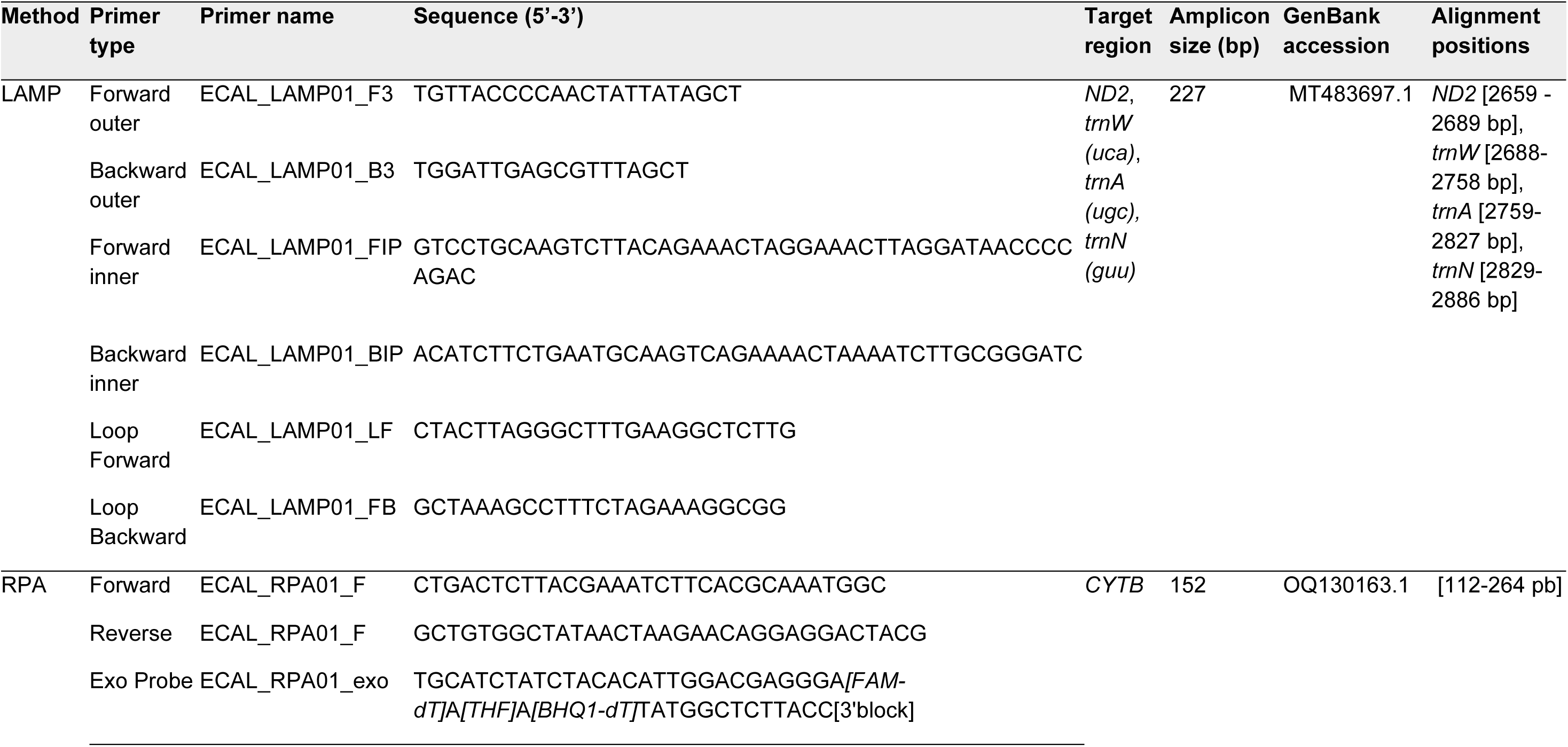
Final primer selection for LAMP and RPA.

### 3.2 Evaluation of metrics for a positive and specific amplification

The TTT metric showed significant difference between target and non-target DNA for both LAMP and RPA (W = 60, p < 0.0001, **Fig. 3a**). Target DNA was detected earlier, with a median TTT of 14.8 min for LAMP and 4.5 min for RPA, compared to 45 min and 31.9 min for the other species, respectively. In contrast for qPCR, we did not observe significant differences in TTT values for the target and non-target species due to a huge variability for non-target species (W = 26, p = 0.7021, **Fig. 3a**). The approximate median reaction time for DNA samples from *E. calamita* is around 72.8 min, equivalent to 25.12 cycles. Looking at EndRFU values, amplifications of the target species were significantly higher than that of non-target species for all methods (W = 0, p < 0.001, **Fig. 3b**).

**Figure 3.**
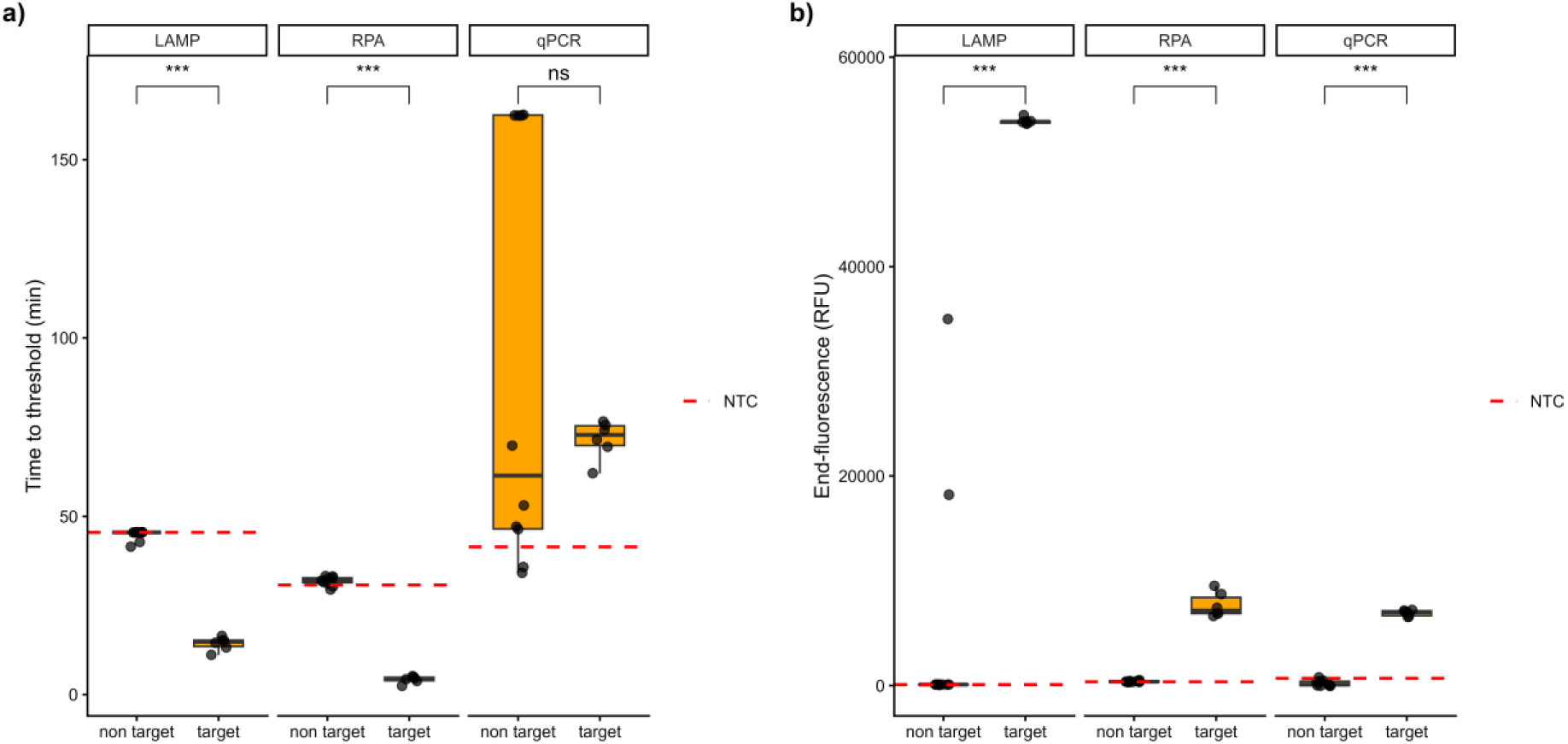
Amplification efficiency of tissue extracts from *Epidalea calamita* versus close related species for LAMP, RPA and qPCR. Each point is the average value of a) Time to threshold values or b) End-point fluorescence values for the three technical replicates. For Mann-Whitney U tests: *** indicates a p-value < 0.001, while ns indicates a non-significant result.

### 3.3 Assay sensitivity

The GLM model explained 79.9% of the observed variability in detection probability. DNA concentration showed the strongest effect on detection probability, by accounting for 72.1% of the total deviance of the model. Initial DNA concentration had a strong positive effect on detection probability (log-odds estimate =1.8863, se = 0.2945, z = 6.405, p < 0.0001). LAMP had the lower detection probability compared to qPCR (log-odds estimate = −2.545, se = 0.755, z = −3.373, p < 0.001). By contrast, RPA probability of detection did not differ significantly from qPCR (p = 0.198). Post-hoc comparisons based on estimated marginal means at the mean DNA concentration showed that qPCR had significantly higher detection odds than LAMP (odds ratio = 12.5, p = 0.002), whereas no significant difference was detected between qPCR and RPA or between RPA and LAMP. The LOD95 values, estimated from the above model for each method, are presented in **Table 2**. The 95% confidence intervals of LAMP and qPCR overlapped by 1.17 pg/ul (18.2% of the narrower interval), those of RPA and qPCR by 5.96 pg/µL (92.8%), and those of RPA and LAMP by 13.89 pg/µL (74.4%, **Table 2**).

**Table 2.** Limit of detection and confidence intervals obtained from LAMP, RPA and qPCR sensitivity results and model predictions.

| **Method** | **LOD95 (pg/ul)** | **CI_inf (pg/ul)** | **CI_sup (pg/ul)** |
|---|---|---|---|
| LAMP | 20.2371710 | 5.5143374 | 155.53867 |
| RPA | 2.5790534 | 0.7076266 | 19.38867 |
| qPCR | 0.9061809 | 0.2496657 | 6.67270 |

### 3.4 Validation on eDNA samples

Almost all eDNA samples coming from sites where the species presence was confirmed by field scientists gave positive amplifications for all three methods. The only exception was a Toulouse biological replicate, which was not amplified by isothermal methods, but for which only one technical replicate was found positive by qPCR. Samples with confirmed absence or uncertain presence all gave negative results. Negative extraction controls and eDNA samples with other amphibians also gave negative results. The two concentrations of positive control yielded positive detections for both qPCR and RPA. For LAMP, the 13.7 pg/μL dilution was positive for one technical replicate out of three (**Fig. 4**).

**Figure 4.**
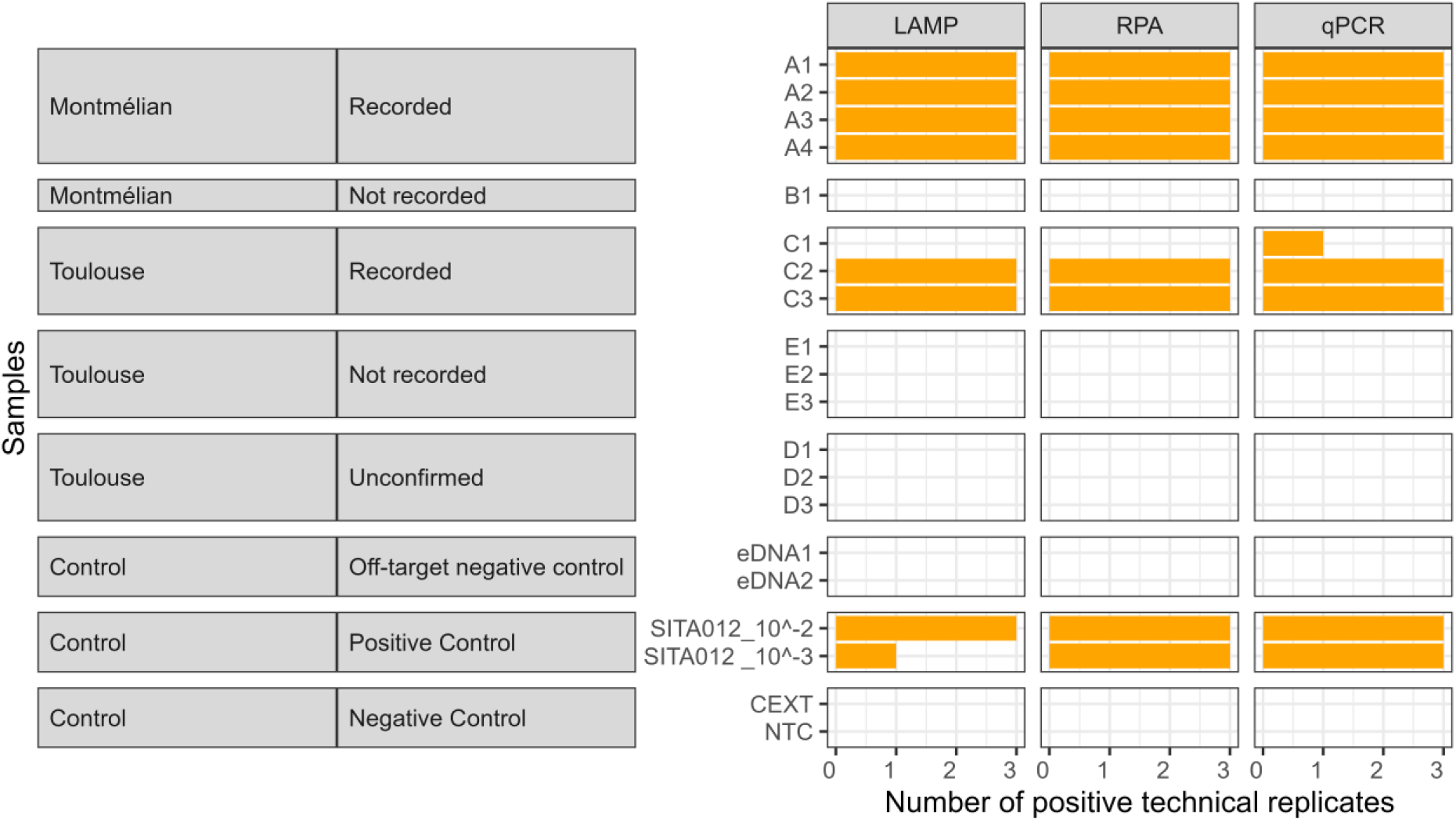
Success rate of amplification methods on eDNA samples. Sample names refer to the sampling sites and replicates presented in figure 2. Recorded: species detected visually or acoustically during the survey; Not recorded: species not detected during the survey; Unconfirmed: reported or suspected presence that could not be reliably confirmed on the day of sampling. The positive control consisted of a tissue sample from *E. calamita*, the same sample used for sensitivity analyses. Negative controls consisted of extraction control (CEXT) and a negative template control (NTC).

## 4 Discussion

In the present study, we explored two promising isothermal amplification methods for eDNA applications with high theoretical reaction speed, potentially high specificity and sensitivity, and minimal equipment requirements. After the design of different sets of LAMP and RPA primers specific to *E. calamita in silico,* our tests allowed identifying the primers and protocols that ensure a clear interpretation of the results (**Table 1**). Specificity tests showed that our LAMP and RPA assays both yielded positive results for the target species, even for genomic DNA samples collected from different populations, individuals, states of development, and tissue types (**Table S1**). We conclude that there is no cross-amplification with *R. temporaria*, *B. bufo*, *B. variegata*, *B. viridis* and *B. spinosus* (**Fig 3**). This demonstrates that our assays are conservative enough to detect *E. calamita* DNA from genetically diverse populations, and that they are specific against sympatric and genetically close related species tested. Validation on eDNA samples further confirmed the capacity of our LAMP and RPA assays to detect with confidence the species’ DNA in complex samples where the species is certified as present by field scientists (**Fig. 4**). We further validated our protocols by conducting additional cross-amplification tests on off-target eDNA samples from ponds harboring other amphibian species (**Table S7**). The absence of amplification in these samples provided further evidence confirming the specificity of the tests (**Fig. 4**). Still, further validation by cross-reactivity assessment may be needed for additional amphibian species that may occur within the distribution range of *E. calamita* not included in our experiment such as *Hyla molleri*, *Pelophylax perezi, Pelobates cultripes, Pelodytes punctatus*, *Discoglossus pictus* as well as urodeles (Fauchier et al. 2017; Sánchez-Montes et al. 2018; Pujol-Buxó et al. 2019).

In parallel, the samples were also analyzed using qPCR to compare the detection performance of the three amplification methods. Because the three assays target distinct mitochondrial markers, the results should be interpreted as a comparison of assay performance rather than a strict methodological comparison. As with LAMP and RPA, we first evaluated the specificity of the established qPCR primers against closely related species **(Fig. 3**; Blom 2018). During these tests, amplification curves were observed in some non-target and negative control samples across all methods (**Fig S2**). We therefore compared the EndRFU and TTT metrics to determine which best discriminate between target and non-target amplification curves. For LAMP and RPA, both TTT and EndRFU values successfully discriminated against the two groups signals with significant difference (**Fig 3**). In contrast, for qPCR, the EndRFU measurement was more effective than the TTT measurement at distinguishing between the groups (**Fig. 3**). This difference may be partly explained by the default automatic threshold settings of our CFX Maestro across all runs. Under these conditions, some atypical amplification curves, usually considered background noise, yielded spurious Cq values in qPCR (**Fig S2a**). As EndRFU provided strong discrimination between target and non-target groups and was consistent for all methods we selected it as our principal detection criterion throughout the study. Based on this criterion, we therefore validate with reliance the specificity of the qPCR primers. Thus, TTT remains informative for defining an appropriate maximum reaction time. This is particularly important when developing rapid detection assays, as extending reaction times may not necessarily improve detection performance and may instead increase the likelihood of non-specific amplification and lead to false positive detection (Zyrina and Antipova 2021). The recommendations for the protocols we use are 30 min for LAMP and 20 min for RPA. Based on our results, reaction times could be limited to 40 min for LAMP and 30 min for RPA. Optimal reaction times are assay and concentration dependent. Determining the optimal reaction is finding the balance between rapidity of the results, sensitivity of the assay, and limitation of false positives.

Sensitivity of LAMP was 10-fold lower than that of qPCR (**Table 2**), and it was also more variable, as visible by the large 95% confidence intervals of LAMP LOD95s. A larger sample size, particularly around the LOD95 region, will most likely help to narrow these confidence intervals and clarify the difference between LAMP and the other methods. In contrast, the RPA assay did not significantly differ from that of the qPCR. In the literature, similar differences between isothermal methods and qPCR have been reported. For example, (Carvalho et al. 2021) reported a ten-fold higher sensitivity of qPCR (0.056 pg /μL) compared to that of LAMP (0.56 pg /μL) for *Dreissena polymorpha* DNA. (Kageyama et al. 2022) also found LAMP to be two orders of magnitude less sensitive than qPCR for grass carp. Other studies have also reported equivalent LODs for RPA and qPCR for *Salmonella spp* (Zhao et al. 2021). However, these observations are not universal: other studies reported comparable or up to 100-fold higher sensitivity of LAMP than qPCR on pathogens or macroorganisms (Jang et al. 2024; Jothinarayanan et al. 2023; Foo et al. 2020). Likewise, RPA was sometimes also found to be more sensitive than qPCR on eDNA samples, when combined with CRISPR-Cas technology (D’Agnese et al. 2023; Wei et al. 2023).

When applied to eDNA samples, isothermal techniques, qPCR and traditional monitoring results showed overall consistency for the detection of *E. calamita*. We expected eDNA methods to fail to detect the species where their presence was uncertain (**Site D, Fig. 2b**). At this site, only one individual had been observed by field scientists nearby the same year, days before the sampling session, and since then, there had been no sightings of singing or egg-laying. In one pond from the Toulouse site, LAMP and RPA failed to detect the target DNA, despite amplification being successful in one qPCR technical replicate. This may be explained by low target DNA concentrations in this sample, so low that they are close to the limit of detection (sample C1, **Fig. 4**). As amplification is performed on heterogeneous DNA samples containing low copy numbers of the target and potentially inhibitory compounds, its efficiency and the probability of detection resulting are affected. A diagnostic sensitivity analysis could be expanded, by amplifying eDNA samples with a more subtle gradient in species abundance and by adding more biological or technical replicates in the analysis to better support decision making when DNA is scarce. Further research could be also done, for example regarding LAMP/RPA *versus* qPCR on inhibitor tolerance, as both have been reported to show greater tolerance to several PCR inhibitors than conventional PCR (Nwe et al. 2024; Singh et al. 2026). A lower sensitivity of isothermal methods applied to eDNA samples may indeed be compensated by a greater stability of detection across environmental conditions, a well-known issue of PCR. These results could therefore provide additional insights into the diagnostic sensitivity of isothermal techniques compared to qPCR in applications involving eDNA. In addition, their reported relative tolerance to inhibitors makes them promising tools for field-based DNA amplification, where extraction conditions are often less controlled than in the laboratory and may yield less purified DNA.

In practice, RPA and LAMP primer design were roughly equivalent but remain today more empirical and labor-intensive than PCR. On one hand, the methodology used for LAMP did not allow us to properly assess primer specificity *in silico.* In fact, the limited availability of sequences from other amphibian species in genomic databases for the selected marker restricted the assessment of primer specificity. Tools such as BLAST (Altschul et al. 1990) could have been combined to further assess primer specificity, although the large number of primers required for LAMP would have made this approach more time-consuming. We also know that specific primer design is more challenging for LAMP compared to RPA or qPCR due to the risk of primer-interactions (Yan et al. 2024; Zyrina and Antipova 2021). In our case, NEB’s LAMP primer design tool greatly facilitated initial primer selection, despite the complexity of the design. For RPA, although relatively few tools have been specifically developed for primer design, dedicated tools such as PrimedRPA (Higgins et al. 2019) are available for this purpose. However, PCR primer design tools such as Primer-BLAST can also be used by adapting the selection criteria to RPA requirements, as in our study following the TwistAmp manual recommendations and the strategy described by (Kellner et al. 2019). A key challenge resides in the specificity of the primers as the literature indicates that RPA is more prone to cross-amplification due to a higher tolerance for mismatches compared to qPCR (Singh et al. 2026; Zou et al. 2020; Hueso et al. 2025). To limit this risk, we complemented primer design with an additional *in silico* alignment of sequences retrieved from NCBI. This allowed us to examine sequence differences in the candidate primer-binding regions and select primers expected to maximize discrimination between the target and non-target species.

In the laboratory, RPA required more pipetting and greater precision compared to the other amplification methods. The reaction uses more enzymes and requires more reagents than LAMP and qPCR. On the other hand, both LAMP and RPA are available in lyophilized form, reducing the number of handling steps since all that is required is to rehydrate the solution and add the primers and the sample. Beyond analytical and technical considerations, the cost per reaction is an important consideration for the practical implementation of these assays. In our study, the estimated cost per reaction of RPA was higher (∼ 3.5 euros/reaction) followed by LAMP (∼1.6 euros/reaction) and qPCR (∼1.2 euros/reaction). This considers consumables only, including the amplification kits, primers, probes, and additional reagents required beyond those supplied in the kits (e.g., dNTPs for RPA and BSA for qPCR).

Overall, although RPA performance indicators must be balanced against practical and financial considerations, the RPA assay proposed in this study emerges as the fastest reaction and was sufficiently sensitive for detection of *E. calamita* on aquatic eDNA samples. Primers and protocols used in this study could therefore be applied more broadly for the detection of this species. Moreover, the methodology we followed is transferable and can be used to develop new isothermal amplification tests for other species. This work also offers other interesting research perspectives. First, sequencing the amplicon products would enable sequence-level validation of positive detections and assessment of occurrence of false-positive detection (Yugovich et al. 2026). Depending on the targeted region and sequencing approach, it could also provide information on amplicon sequence diversity. Second, it would also allow the analysis to go beyond simple target detection, would open the way to downstream applications like metabarcoding of eDNA (Plewnia et al. 2025). Finally, perhaps the most exciting perspective lies in the ease of deployment of isothermal amplifications in the field. Besides providing quick results, LAMP and RPA rely only on simple constant temperature heating systems and may tolerate certain inhibitors present in partially purified DNA extracts obtained with simplified extraction protocols. Although fluorescence-based detection, as used in our study, still requires dedicated instrumentation, alternative readout methods, such as colorimetric detection or lateral-flow assays, could further reduce equipment requirements. Combined with appropriate field-based workflow for sample preparation and portable heating systems, these approaches could ultimately enable rapid *in situ* eDNA detection. All these features make these nucleic acid amplification techniques particularly suitable for these applications.

## Authors’ contributions

A.C., A.B., L.Z., A.I., A.G., P.T. and E.B. conceived and designed the study. A.C. performed the acquisition and analysis of the data, with the help of A.G. and F.B.. A.C., A.B., A.I. and L.Z. interpreted the results. A.C. wrote the first version of the manuscript, with the help of A.B., A.G., A.I. and L.Z., and all authors contributed to its revision. A.C., A.B., A.I., L.Z. and T.M. obtained the funding. A.I., A.B. and L.Z. supervised the work.

## Data archiving statement

Raw amplification data, processed data, metadata and scripts to produce statistical analyses and graphical representations will be publicly available upon acceptance in the InDoRES data repository (https://doi.org/10.48579/PRO/5UKFZN).

## Acknowledgments

We acknowledge the French “Collection CEFE - EPHE de l’équipe Biogéographie et Ecologie des Vertébrés de Montpellier (collection BEV)”, and in particular Philippe Geniez for providing tissue samples. We thank Jérémie Hahn (TEREO, Montmélian, France) and the team at Nature en Occitanie (Toulouse, France) for their assistance with field site selection and survey validation. We also thank Pablo Tedesco, the Argaly team, and Nature en Occitanie for their help with sampling. We thank Claire Sevin and Elisabeth Guichard from the Office National des Forêts (agence Ile-de-France Ouest) for providing environmental samples. The authors gratefully acknowledge all funding bodies for their financial support.

