## Supplementary figures for "Rapid environmental DNA detection with isothermal amplification methods: a comparison with the real-time quantitative PCR in the natterjack toad (*Epidalea calamita*)"

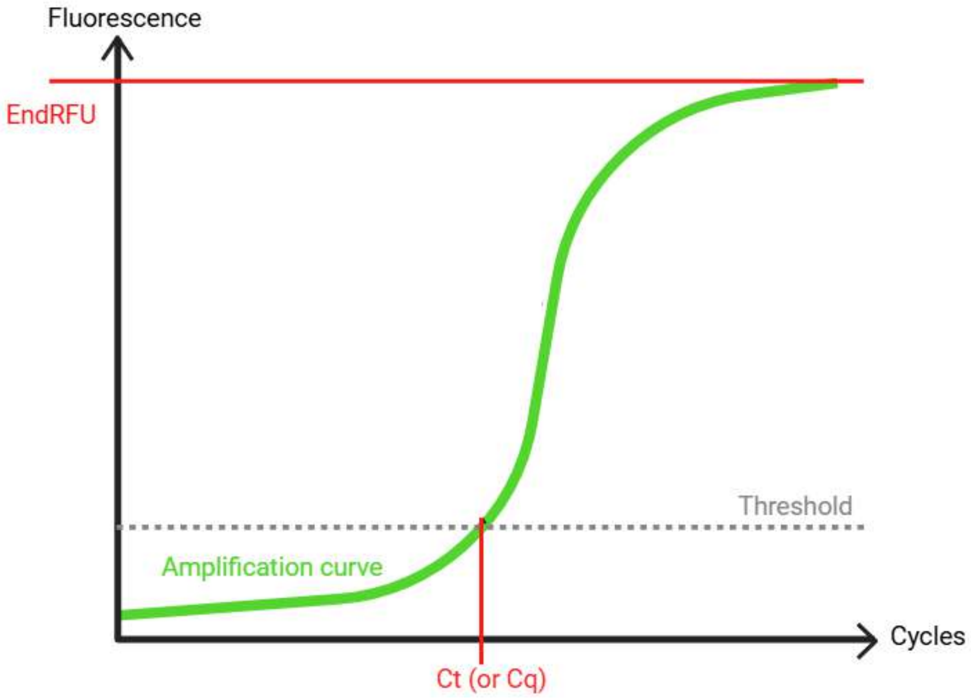


**Figure S1: Theoretical graph of amplification curves with explanation of the threshold cycle (Ct) (or quantification cycle (Cq)) and end point fluorescence units (EndRFU) values.** The Ct value corresponds to the point where the curve intersects with the defined baseline fluorescence threshold. The EndRFU value corresponds to the RFU value of the curve at a specific point in time defined as the end of the reaction.

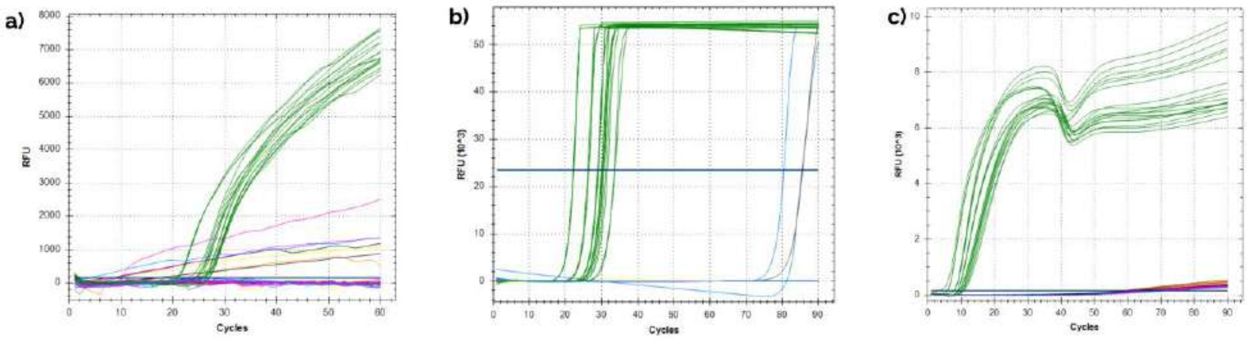


**Figure S2: Amplification curves of primer specificity tests for a) qPCR b) RPA and c) LAMP.** For each amplification, three replicates of each extract have been realized. DNA extracts from *E.calamita* are in green (x6 individuals), *B.variegata* in yellow (x2), *B.bufo* in brown (x2), *B.spinosus* in pink (x3), *B.viridis* in blue (x2) and *R.temporaria* in orange (x1).
