## Supplementary tables for "Rapid environmental DNA detection with isothermal amplification methods: a comparison with the real-time quantitative PCR in the natterjack toad (*Epidalea calamita*)"

Table S1: LAMP Primer Design default parameters used in our study

| | | |
|---|---|---|
| **Sorting Rule** | | Easy |
| **Reset parameters** | | Normal default parameters |
| **Reaction Conditions** | $Na^+$ concentration | 50 |
| | $Mg^{++}$ concentration | 8 |
| **Lengths (min/max)** | F1c/B1c | 20-22 |
| | F2/B2 | 18-20 |
| | F3/B3 | 18-20 |
| | LF/LB | 15-25 |
| **$T_m$ (min/max)** | F1c/B1c | 64-66 |
| | F2/B2 | 59-61 |
| | F3/B3 | 59-61 |
| | LF/LB | 64-66 |
| **% GC** | Min/Max | 40-65 |
| | Min/Max (Loop) | 40-65 |
| **ΔG threshold** | 5' Stability | -4 |
| | 3' Stability | -4 |
| | 3' Stability (Loop) | -2 |
| | Dimer check | -2,5 |
| | Dimer check (Loop) | -3,5 |
| **Distances (min/max)** | (F2-B2) | 120-160 |
| | Loop (F1c-F2) | 40-60 |
| | F2-F3 | 0-60 |
| | F1c-B1c | 0-100 |
| **Limits** | F1c/B1c | 3 |
| | F2/B2 | 10 |
| | F3/B3 | 3 |
| | LF/LB | 10 |
| | Sets | 1000 |

Table S2: List of LAMP primer set selected for in vitro analysis.

| **LAMP Primer set (ID; sequence)** | **Target region** |
|---|---|
| **ECal_LAMP01_F3 ;** GTTAAAGAATCCTATTCTAAGCCAA | L-rRNA |
| **ECal_LAMP01_B3 ;** AGGGTTTTAATCGTTGAACAA | |

| | |
|---|---|
| **ECal_LAMP01_FIP ;** GCTGTTATCCCTAGGGTAACTTGGGCATCTAAGCATCAACACATT | |
| **ECal_LAMP01_BIP ;** CTATCGACAAGTGGGTTTACGACACGAACCATTAGTAGCGG | |
| **ECal_LAMP01_LF ;** TGCTCAAATGTATTGGGTCAATGGA | |
| **ECal_LAMP01_LB ;** TGTTGGATCAGGGTATCCTAGTGG | |
| **ECal_LAMP02_F3 ;** TGTTACCCCAACTATTATAGCT | NADH dehydrogenase subunit 2 + trnW(uca) + trnA(ugc) +trnN(guu) |
| **ECal_LAMP02_B3 ;** TGGATTGAGCGTTTAGCT | |
| **ECal_LAMP02_FIP ;** GTCCTGCAAGTCTTACAGAAACTAGGAAACTTAGGATAACCCCAGAC | |
| **ECal_LAMP02_BIP ;** ACATCTTCTGAATGCAAGTCAGAAAACTAAAATCTTGCGGGATC | |
| **ECal_LAMP02_LF ;** CTACTTAGGGCTTTGAAGGCTCTTG | |
| **ECal_LAMP02_LB ;** GCTAAAGCCTTTCTAGAAAGGCGG | |
| **ECal_LAMP03_F3 ;** TGGACAATCGAATAACCACC | cytochrome c oxidase subunit II |
| **ECal_LAMP03_B3 ;** GCAAATCTCGGAACACTGT | |
| **ECal_LAMP03_FIP ;** AAGACGGACAGCTCATGAGTGCCAGTAGGTATAACAACCCG | |

| | |
|---|---|
| **ECal_LAMP03_BIP ;** GGCGTAAAATTAGACGCAATTCCTAGTAAACGCCAGGTCGG | |
| **ECal_LAMP03_LF ;** ACATTCTTCCTGCAGTAATCAGGG | |
| **ECal_LAMP03_LB ;** AGGACGACTAAATCAGACCTCGT | |
| **ECal_LAMP04_F3 ;** GATATTCGAAAAATAGGGGGC | NADH dehydrogenase subunit 5 |
| **ECal_LAMP04_B3 ;** CGGGTGTTTTATTGATGCA | |
| **ECal_LAMP04_FIP ;** TAGAAAAAAATCCTGCGAGGTAGGGCCATTTACAACAACATGCGT | |
| **ECal_LAMP04_BIP ;** CAATCAACACCTCCAACGTTAATGCTTACGGAGGCTGTAGACG | |
| **ECal_LAMP04_LF ;** GTTCCTATAAGGGCCAGACTTCCT | |
| **ECal_LAMP04_LB ;** CCCTTACACTGACTATCATTGCCAC | |
| **ECal_LAMP05_F3;** ACACGCCTGTTATCAAGAA | 12S rRNA |
| **ECal_LAMP05_B3;** TTATGGACAGGCTCCTCTA | |
| **ECal_LAMP05_FIP;** ATAGTGGGGTATCTAATCCCAGTTTCTAAAGTCGCTCTAACACAATC | |
| **ECal_LAMP05_BIP;** CTCAAACCGCCAGGGAACTAGGGATATGGGGTACCGTC | |

| |
|---|
| **ECal_LAMP05_LF;** CCTAGCTGTCGTGAGTTCAAGTT |
| **ECal_LAMP05_LB;** CGAGCAAAGCTTAAAACCCAAAGG |

Table S3: Table of custom parameters in NCBI Primer-BLAST for E. calamita RPA primer templates.

| **Parameters** | **Values** |
|---|---|
| PCR product size | [80;200] |
| Number of primers to return | 100 |
| Primer melting temperature | [50;100] |
| Max Tm difference | 1 |
| Database | nt |
| Organism | Epidalea calamita (taxid: 30326) |
| Advanced parameters | Values |
| Primer size | [30 ;35] |
| Primer GC content (%) | [40 ;70] |

Table S4: Table of DNA sequences from potential false positives (closely related species of Epidalea calamita) retrieved from NCBI.

| **Species** | **NCBI accession number** | **Target** |
|---|---|---|
| *Hyla meridionalis* | MK511412.1 | COI |
| *Hyla meridionalis* | MK013993.1 | COI |
| *Hyla molleri* | JN801026.1 | COI |
| *Hyla molleri* | JN801025.1 | COI |
| *Hyla arborea* | MK511411.1 | COI |
| *Hyla arborea* | MK013988.1 | COI |
| *Hyla sarda* | MK013998.1 | COI |
| *Lithobates catesbeianus* | OR150031.1 | COI |
| *Aquarana catesbeiana* | PP543528.1 | COI |
| *Pelophylax bedriagae* | MT881779.1 | COI |
| *Pelophylax bedriagae* | MT881778.1 | COI |
| *Pelophylax esculentus* | KP697836.1 | COI |

| | | |
|---|---|---|
| *Pelophylax grafi* | KT879323.1 | COI |
| *Pelophylax lessonae* | MN993185.1 | COI |
| *Pelophylax perezi* | KT879335.1 | COI |
| *Pelophylax ridibundus* | MN993187.1 | COI |
| *Rana saharica* | HM363049.1 | COI |
| *Rana arvalis* | MT872666.1 | COI |
| *Rana dalmatina* | MT876986.1 | COI |
| *Rana dalmatina* | MT876988.1 | COI |
| *Rana pyrenaica* | KC977251.1 | COI |
| *Rana temporaria* | KT074473.1 | COI |
| *Rana temporaria* | MT876989.1 | COI |
| *Alytes obstetricans* | JQ626705.1 | COI |
| *Discoglossus montalentii* | JQ626690.1 | COI |
| *Discoglossus montalentii* | JQ626689.1 | COI |
| *Discoglossus pictus* | JQ626698.1 | COI |
| *Discoglossus pictus* | JQ626697.1 | COI |
| *Discoglossus sardus* | JQ626696.1 | COI |
| *Discoglossus sardus* | JQ626695.1 | COI |
| *Bombina variegata* | MN993093.1 | COI |
| *Bombina variegata* | MN993092.1 | COI |
| *Bombina variegata* | MN993091.1 | COI |
| *Bombina bombina* | EU115993.1 | COI |
| *Bombina bombina* | JX893172.1 | COI |
| *Bufo bufo* | MH734357.1 | COI |
| *Bufo bufo* | MK013982.1 | COI |
| *Bufo spinosus* | MK511409.1 | COI |
| *Bufo spinosus* | MK511408.1 | COI |
| *Bufotes viridis* | MN993105.1 | COI |
| *Bufotes viridis* | MN993104.1 | COI |
| *Pelobates fuscus* | MN993177.1 | COI |
| *Pelobates fuscus* | MN993176.1 | COI |
| *Pelobates cultripes* | AJ871086.1 | COI |
| *Pelodytes punctatus* | MK014017.1 | COI |
| *Xenopus laevis* | OQ261767.1 | COI |
| *Xenopus laevis* | OQ318522.1 | COI |
| *Xenopus laevis* | OQ318521.1 | COI |
| *Lissotriton vulgaris* | MK014006.1 | COI |
| *Lissotriton vulgaris* | MN993150.1 | COI |
| *Lissotriton helveticus* | JF812747.1 | COI |

| | | |
|---|---|---|
| *Lissotriton helveticus* | JF812746.1 | COI |
| *Triturus cristatus* | KP697896.1 | COI |
| *Triturus cristatus* | MN993235.1 | COI |
| *Triturus carnifex* | MN993232.1 | COI |
| *Triturus carnifex* | MN993230.1 | COI |
| *Ichthyosaura alpestris* | MN993134.1 | COI |
| *Ichthyosaura alpestris* | MN993133.1 | COI |
| *Triturus marmoratus* | JN379828.1 | COI |
| *Triturus marmoratus* | JN379827.1 | COI |
| *Salamandra salamandra* | MN993223.1 | COI |
| *Salamandra salamandra* | MN993222.1 | COI |
| *Salamandra atra* | MN993218.1 | COI |
| *Salamandra atra* | MN993217.1 | COI |
| *Salamandra corsica* | MK014026.1 | COI |
| *Salamandra corsica* | KT365355.1 | COI |
| *Euproctus montanus* | DQ517820.1 | COI |
| *Euproctus montanus* | JX626594.1 | COI |
| *Calotriton asper* | EU880307.1 | COI |
| *Speleomantes strinatii* | MK014000.1 | COI |
| *Homo sapiens* | KC750830.1 | COI |
| *Hyla meridionalis* | MK902650.1 | Cytb |
| *Pelophylax ridibundus* | OP344941.1 | Cytb |
| *Hyla arborea* | OP690620.1 | Cytb |
| *Hyla sarda* | MW570679.1 | Cytb |
| *Lithobates catesbeianus* | OP414972.1 | Cytb |
| *Pelophylax bedriagae* | AB981660.1 | Cytb |
| *Pelophylax lessonae* | MT737359.1 | Cytb |
| *Pelophylax perezi* | MF667576.1 | Cytb |
| *Pelophylax ridibundus* | MG575226.1 | Cytb |
| *Rana saharica* | GU799108.1 | Cytb |
| *Rana arvalis* | OQ130159.1 | Cytb |
| *Rana dalmatina* | OQ130165.1 | Cytb |
| *Rana pyrenaica* | AY147971.1 | Cytb |
| *Rana pyrenaica* | AY043059.1 | Cytb |
| *Rana temporaria* | OQ130162.1 | Cytb |
| *Alytes obstetricans* | AY151783.1 | Cytb |
| *Discoglossus montalentii* | KC342987.1 | Cytb |
| *Discoglossus pictus* | MH142696.1 | Cytb |
| *Discoglossus sardus* | MT569387.1 | Cytb |

| | | |
|---|---|---|
| *Bombina variegata* | MW296356.1 | Cytb |
| *Bombina bombina* | MN784161.1 | Cytb |
| *Bufo bufo* | AB159262.1 | Cytb |
| *Bufo spinosus* | JN647460.1 | Cytb |
| *Bufotes viridis* | OQ130164.1 | Cytb |
| *Pelobates fuscus* | MK902549.1 | Cytb |
| *Pelobates cultripes* | GU983095.1 | Cytb |
| *Pelodytes punctatus* | JF272534.1 | Cytb |
| *Xenopus laevis* | KT587048.1 | Cytb |
| *Lissotriton vulgaris* | DQ821272.1 | Cytb |
| *Lissotriton helveticus* | DQ821239.1 | Cytb |
| *Triturus cristatus* | EU030898.1 | Cytb |
| *Triturus carnifex* | EU030818.1 | Cytb |
| *Ichthyosaura alpestris* | KY911419.1 | Cytb |
| *Triturus marmoratus* | EU030900.1 | Cytb |
| *Salamandra salamandra* | MN437482.1 | Cytb |
| *Salamandra atra* | MG968419.1 | Cytb |
| *Salamandra corsica* | KT365391.1 | Cytb |
| *Salamandra lanzai* | AY928613.1 | Cytb |
| *Euproctus montanus* | JX626788.1 | Cytb |
| *Calotriton asper* | MT498349.1 | Cytb |
| *Speleomantes strinatii* | OR353875.1 | Cytb |
| *Homo sapiens* | JN034136.1 | Cytb |

Supplementary tables

Table S5: List of RPA primers selected for in vitro analysis.

| Primer ID | Sequence | Primer ID | Sequence | Exo probe ID | Sequence |
|---|---|---|---|---|---|
| ECal_RPA_Cytb_F1 | CTGACTCTTACGAAATCTTCACGCAAATGGC | ECal_RPA_Cytb_R1 | GCTGTGGCTATAACTAAGAACAGGAGGACTACG | ECal_RPA_Cytb_Exo | TGCATCTATCTACACATTGGACGAGGGA[FAM-dT]A[THF]A[BHQ1-dT]TATGGCTCTTACC[3'block] |
| ECal_RPA_CytB_F2 | GCTGACTCTTACGAAATCTTCACGCAAATGG | ECal_RPA_Cytb_R2 | AATGCTGTGGCTATAACTAAGAACAGGAGGACTA | ECal_RPA_Cytb_Exo | idem |
| ECal_RPA_CytB_F3 | GGCTGACTCTTACGAAATCTTCACGCAAATG | ECal_RPA_Cytb_R2 | GGCTATAACTAAGAACAGGAGGACTACGCCAATA | ECal_RPA_Cytb_Exo | idem |
| ECal_RPA_COI_F1 | TTTACTCCTTCTGGCCTCCGCTGGGGTAGAG | ECal_RPA_COI_R1 | TTGACGGGGGTTTTATGTTCAGGGTGGTGGTAA | ECal_RPA_COI_Exo | CTGACTATTTTTTCCCTTCATCTTGCAGG[FAM-dT]G[THF]A[BHQ1-dT]CCTCTATTCTAGGG[3'block] |
| ECal_RPA_COI_F2 | TTACTCCTTCTGGCCTCCGCTGGGGTAGAG | ECal_RPA_COI_R2 | ATTGACGGGGGTTTTATGTTCAGGGTGGTGGT | ECal_RPA_COI_Exo | idem |
| ECal_RPA_COI_F3 | TTTTACTCCTTCTGGCCTCCGCTGGGGTAGAG | ECal_RPA_COI_R3 | TGACGGGGGTTTTATGTTCAGGGTGGTGGT | ECal_RPA_COI_Exo | idem |

Supplementary tables

Table S6: List of amphibian's tissues samples used in the study (part1).

| Supplier | Sample ID | N_BEV | Origin | Family | Genus_species | taxon | J | M | Year | ISO |
|---|---|---|---|---|---|---|---|---|---|---|
| BEV collection | SITA001 | photo PAC | field | Bombinatoridae | Bombina variegata | Bombina variegata scabra | 9 | 8 | 2022 | MKD |
| | SITA002 | photo PAC | field | Bombinatoridae | Bombina variegata | Bombina variegata scabra | 9 | 8 | 2022 | MKD |
| | SITA003 | voucher | field | Bufonidae | Bufo bufo | Bufo bufo bufo | 7 | 6 | 2015 | FRA |
| | SITA004 | voucher | field | Bufonidae | Bufo bufo | Bufo bufo bufo | 29 | 2 | 2024 | FRA |
| | SITA005 | BEV.15490 | field | Bufonidae | Bufo spinosus | Bufo spinosus | 20 | 6 | 2020 | FRA |
| | SITA006 | BEV.15500 | field | Bufonidae | Bufo spinosus | Bufo spinosus | 20 | 6 | 2020 | FRA |
| | SITA007 | voucher | field | Bufonidae | Bufo spinosus | Bufo spinosus | 3 | 5 | 2022 | FRA |
| | SITA008 | voucher | field | Bufonidae | Bufotes viridis | Bufotes viridis viridis | 24 | 3 | 2019 | GRC |
| | SITA009 | BEV.16039 | field | Bufonidae | Bufotes viridis | Bufotes viridis sitibundus | 4 | 4 | 2023 | JOR |
| | SITA010 | photo PAC | field | Bufonidae | Epidalea calamita | Epidalea calamita | 19 | 5 | 2011 | FRA |
| | SITA011 | BEV.15853 | jar | Bufonidae | Epidalea calamita | Epidalea calamita | 22 | 4 | 2022 | ESP |
| | SITA012 | BEV.15181 | field | Bufonidae | Epidalea calamita | Epidalea calamita | 24 | 4 | 2018 | FRA |
| | SITA013 | BEV.15182 | field | Bufonidae | Epidalea calamita | Epidalea calamita | 24 | 4 | 2018 | FRA |
| | SITA014 | BEV.16541 | field | Bufonidae | Epidalea calamita | Epidalea calamita | 9 | 5 | 2001 | FRA |
| | SITA015 | BEV.16542 | field | Bufonidae | Epidalea calamita | Epidalea calamita | 9 | 5 | 2001 | FRA |
| | SITA017 | BEV.15672 | field | Ranidae | Rana temporaria | Rana temporaria | 1 | 5 | 2021 | FRA |

Table S6: List of amphibian's tissues samples used in the study (part2).

Supplementary tables

| Dept | Collection_site | Latitude | Longitude | Alt | Precision_m | Collectors |
|---|---|---|---|---|---|---|
| | Zovich (= Zovikj), at the bridge | 41,11887 | 21,71709 | 645 | 30 | Pierre-André Crochet, Dragan Arsovski |
| | Zovich (= Zovikj), at the bridge | 41,11887 | 21,71709 | 645 | 30 | Pierre-André Crochet, Dragan Arsovski |
| 38 | Theys, 500m SSE of Cul de Pet | 45,2906 | 6,0409 | 1840 | | P.-A. Crochet |
| 74 | Commune of Doussard, D909A road just south of Glière beach, east side of Lake Annecy, wildlife crossing point | 45,7977 | 6,2316 | 450 | 25 | Claude Miaud |
| 66 | Jujols, hillside reservoir of Jujols, Serrat de la Couytive | 42,57647 | 2,28787 | 1170 | 30 | Pierre-André Crochet & Paul Doniol-Valcroze |
| 66 | Jujols, hillside reservoir of Jujols, Serrat de la Couytive | 42,57647 | 2,28787 | 1170 | 30 | Pierre-André Crochet & Paul Doniol-Valcroze |
| 66 | D3 road, bend on right when descending, 270 m east of Coustouges | 42,36765 | 2,6539 | 753 | 10 | P.-A. Crochet, Philippe Geniez, Julien Renoult |
| Crete | Crete, 250m NE of Kaliviani | 35,51436 | 23,61057 | 26 | 20 | Pierre-André Crochet |
| | Jordan, at 31.8627 / 35.6801 | 31,8627 | 35,6801 | -125 | | Paul Doniol-Valcroze |
| 42 | Sainte-Foy-Saint-Sulpice, Étang des Marquants, access path near house | | | | | P.-A. Crochet, G. Dumont |
| | Hotel la Escondida, Penaguila (parking) | 38,67731 | -0,38182 | | 20 | Pierre-André Crochet, Jérémie Souchet |
| 83 | Plaine des Maures, 4 km NE of Les Mayons | 43,336 | 6,39813 | 110 | 3 | P.-A. Crochet |
| 83 | Plaine des Maures, 4 km NE of Les Mayons | 43,336 | 6,39813 | 110 | 3 | P.-A. Crochet |
| 76 | Port du Havre, Z.II | | | | | Claude Miaud & Christophe Eggert |
| 76 | Port du Havre, Z.II | | | | | Claude Miaud & Christophe Eggert |
| 73 | Savoie, La Table, Les Etelles, bog | 45,4719 | 6,1535 | 720 | | Claude Miaud |

Table S6: List of amphibian's tissues samples used in the study (part3).

| Sample_type | Owner | Box | Barcode | Storage | Extraction_kit | Extraction_date | Concentration of DNA extracts (ng/ul) |
|---|---|---|---|---|---|---|---|
| adult male (buccal swab) | PAC | EPHE-BEV-P170 | CEFE01-162 | Cold room -20°C | NucleoSpin® Tissue kit (Machery-Nagel) | 04/02/2025 | 8,17 |

| | | | | | | | |
|---|---|---|---|---|---|---|---|
| adult female (buccal swab) | PAC | EPHE-BEV-P171 | CEFE01-163 | Cold room -20°C | NucleoSpin® Tissue kit (Machery-Nagel) | 04/02/2025 | 19,05 |
| small larva | PAC | EPHE-BEV-P117 | CEFE01-115 | Cold room -20°C | NucleoSpin® Tissue kit (Machery-Nagel) | 04/02/2025 | 14 |
| adult buccal swab collected during movement (wildlife crossing CMR study) (photos available) | CMi | EPHE-BEV-P187 | CEFE01-179 | Cold room -20°C | NucleoSpin® Tissue kit (Machery-Nagel) | 04/02/2025 | 0,926 |
| 1 metamorph | PAC | EPHE-BEV-P155 | CEFE01-147 | Cold room -20°C | NucleoSpin® Tissue kit (Machery-Nagel) | 04/02/2025 | 12,9 |
| 1 tadpole, hind legs | PAC | EPHE-BEV-P155 | CEFE01-147 | Cold room -20°C | NucleoSpin® Tissue kit (Machery-Nagel) | 04/02/2025 | 39,5 |
| adult tissue crushed on road | PAC | EPHE-BEV-P168 | CEFE01-160 | Cold room -20°C | NucleoSpin® Tissue kit (Machery-Nagel) | 04/02/2025 | 114 |
| small tadpole | PAC | EPHE-BEV-P150 | CEFE01-142 | Cold room -20°C | NucleoSpin® Tissue kit (Machery-Nagel) | 04/02/2025 | 16,15 |
| small tadpole (tail cut for DNA) | PAC | EPHE-BEV-P179 | CEFE01-171 | Cold room -20°C | NucleoSpin® Tissue kit (Machery-Nagel) | 04/02/2025 | 6,53 |
| adult | PAC | EPHE-BEV-P045 | CEFE01-45 | Cold room -20°C | NucleoSpin® Tissue kit (Machery-Nagel) | 04/02/2025 | 24,6 |
| adult found dead | PAC | EPHE-BEV-P169 | CEFE01-161 | Cold room -20°C | NucleoSpin® Tissue kit (Machery-Nagel) | 04/02/2025 | 21,2 |
| larva in roadside ditch near vineyard | PAC | | | | NucleoSpin® Tissue kit (Machery-Nagel) | 04/02/2025 | 13,65 |
| larva in roadside ditch near vineyard | PAC | | | | NucleoSpin® Tissue kit (Machery-Nagel) | 04/02/2025 | 16,35 |
| large juvenile | CMi | | | | NucleoSpin® Tissue kit (Machery-Nagel) | 04/02/2025 | 13,3 |
| subadult | CMi | | | | NucleoSpin® Tissue kit (Machery-Nagel) | 04/02/2025 | 58,9 |
| many medium tadpoles captured in situ | CMi | | | | NucleoSpin® Tissue kit (Machery-Nagel) | 04/02/2025 | 70,8 |

Table S7 : Amphibians taxa dataset recovered from Argaly metabarcoding analysis on the two eDNA samples from Marly Forest

Supplementary tables

| Scientific_name | Phylum_name | Class_name | Order_name | Family_name | Genus_name | Species_name | Taxid | eDNA1 | eDNA2 |
|---|---|---|---|---|---|---|---|---|---|
| Alytes obstetricans | Chordata | Amphibia | Anura | Alytidae | Alytes | Alytes obstetricans | 8443 | 0 | 2993 |
| Bufo bufo | Chordata | Amphibia | Anura | Bufonidae | Bufo | Bufo bufo | 8384 | 0 | 81426 |
| Ichthyosaura alpestris | Chordata | Amphibia | Caudata | Salamandridae | Ichthyosaura | Ichthyosaura alpestris | 54263 | 1056 | 0 |
| Lissotriton helveticus | Chordata | Amphibia | Caudata | Salamandridae | Lissotriton | Lissotriton helveticus | 256425 | 1746 | 0 |
| Rana dalmatina | Chordata | Amphibia | Anura | Ranidae | Rana | Rana dalmatina | 51331 | 12358 | 8178 |
| Rana temporaria | Chordata | Amphibia | Anura | Ranidae | Rana | Rana temporaria | 8407 | 3796 | 0 |
| Salamandra | Chordata | Amphibia | Caudata | Salamandridae | Salamandra | NA | 57569 | 424 | 0 |
| Triturus cristatus | Chordata | Amphibia | Caudata | Salamandridae | Triturus | Triturus cristatus | 8323 | 115 | 0 |

Table S8 : Automatic baseline threshold and EndRFU cutoff for all runs.

| Runs | Baseline and cutoff | LAMP | RPA | qPCR |
|---|---|---|---|---|
| Specificity | Baseline threshold | 23491,75 | 150,7 | 159,28 |
| Sensitivity_Run1 | Baseline threshold | 10285,46 | 236,85 | 104,59 |
| Sensitivity_Run2 | Baseline threshold | 9556,64 | 269,94 | 175,63 |
| Sensitivity_Run3 | Baseline threshold | 7512,36 | 176,24 | 105,97 |
| eDNA | Baseline threshold | 3488,11 | 92,97 | 37,56 |
| Specificity | End-RFU cut-off | 5581 | 1301 | 1389 |
| Sensitivity_Run1 | End-RFU cut-off | 14652 | 1178 | 2562 |
| Sensitivity_Run2 | End-RFU cut-off | 5872 | 1250 | 2829 |
| Sensitivity_Run3 | End-RFU cut-off | 6524 | 885 | 1291 |
| eDNA | End-RFU cut-off | 3964 | 592 | 439 |

Supplementary tables
